# Investigating the role of axonal localization of sex peptide receptor in post-mating responses of female *Drosophila melanogaster*

**DOI:** 10.1101/2024.08.20.608772

**Authors:** Tianmu Zhang, Xinyue Zhou, Woo Jae Kim

## Abstract

This study elucidates the molecular mechanisms underlying the axonal localization of the sex peptide receptor, a pivotal G-protein coupled receptor in the *Drosophila melanogaster* post-mating response cascade. Utilizing transgenic expression, neuronal labeling, and bioinformatics analyses, we demonstrate that the N-terminal domain of SPR is indispensable for its axonal targeting in both *Drosophila* larval ventral nerve cord neurons and mouse hippocampal neurons. Deletion of the N-terminal domain abolished axonal localization, highlighting its critical role in this process. Intriguingly, the C-terminal domain of SPR appears to play a subordinate role in axonal targeting. Bioinformatical analysis revealed a striking homology between the N-terminus of SPR and the Broad-complex, Tramtrack, and Bric-a-brac/poxvirus and zinc finger family of proteins. The BTB domain, a conserved protein-protein interaction domain within this family, is implicated in diverse cellular processes and axonal targeting. Further investigation into the role of the BTB domain-like region in SPR could provide valuable insights into the molecular underpinnings of axonal targeting and post-mating responses in *Drosophila*. This research contributes to our understanding of the intricate mechanisms governing GPCR localization and function in the context of reproductive biology and neuronal signaling.

## INTRODUCTION

In the model organism *Drosophila melanogaster*, the sex peptide (SP) and its receptor (SPR) are key components of the post-mating response cascade. The SP, a peptide hormone produced in the male reproductive tract, is transferred to the female during copulation and binds to the SPR, which is expressed in the female reproductive tract. This molecular dialogue between sexes is crucial for mediating a variety of physiological and behavioral responses in the female, including sperm storage, ovulation, egg-laying patterns, and changes in mating preferences. The SP-SPR system in *Drosophila* serves as an elegant model for understanding the molecular basis of post-mating phenomena and the evolution of reproductive strategies (Wolfner 2002; Chapman et al. 2003; Kubli 2003).

The SPR in *Drosophila melanogaster* is a G-protein coupled receptor (GPCR) that plays a central role in the post-mating responses of female fruit flies. This receptor is uniquely positioned to mediate the molecular functions of the sex peptide. Females lacking SPR either entirely or only in the nervous system fail to respond to the SP and continue to exhibit virgin behaviors even after mating (Yapici et al. 2008). SPR is expressed in the female’s reproductive tract and central nervous system, and its behavioral functions map to neurons that also express the *fruitless* (*fru*) gene, a key determinant of sex-specific reproductive behavior (Häsemeyer et al. 2009; Yang et al. 2009; Vandersmissen et al. 2013; Feng et al. 2014; Oh et al. 2014; Rezával et al. 2014; Avila et al. 2015; Tsuda et al. 2015; Hussain et al. 2016; Yang et al. 2023). Following mating, the transfer of sex peptide from the male triggers SPR activation in these regions, initiating a cascade of post-mating responses (PMR).

Molecularly, the SPR is characterized by its seven-transmembrane domain structure, which is typical of GPCRs. Upon binding of the SP, the SPR undergoes a conformational change that activates intracellular signaling pathways (Yapici et al. 2008). GPCRs represent a pivotal class of membrane-bound signaling proteins that mediate a myriad of cellular responses to extracellular stimuli (Hauser et al. 2006; Caers et al. 2012; Audsley and Down 2015). The precise spatial localization of GPCRs within the cell membrane is a critical determinant of their functional efficacy and signaling specificity. Recent advances in molecular and cell biology have illuminated diverse mechanisms underlying GPCR targeting and trafficking, including receptor oligomerization, interaction with cytoskeletal elements, and the involvement of lipid rafts. Additionally, the post-translational modification of GPCRs, such as glycosylation and phosphorylation, plays a pivotal role in modulating their intracellular trafficking and membrane compartmentalization. Understanding the intricate processes governing GPCR localization is fundamental to deciphering their signaling pathways and developing targeted therapeutic interventions (Stefan et al. 1998; BOIVIN et al. 2008; Hanlon and Andrew 2015).

In an earlier study, the axonal localization of SPR was identified utilizing a panel of specific anti-SPR antibodies (Fig. 1A-B) (Yang et al. 2009). A significant subset of GPCRs exhibits a polarized distribution across the neuronal plasma membrane, a feature that is crucial in shaping the physiological responses elicited by their activation. While recent investigations have delineated various determinants of GPCR targeting to dendrites, the molecular mechanisms underpinning the axonal targeting and subcellular positioning of these receptors have yet to be fully explored (Lasiecka et al. 2008). In the present study, we examine the mechanisms of axonal localization for SPR in an *in vivo Drosophila* model and a vertebrate hippocampal neuronal model.

**Figure 1.**
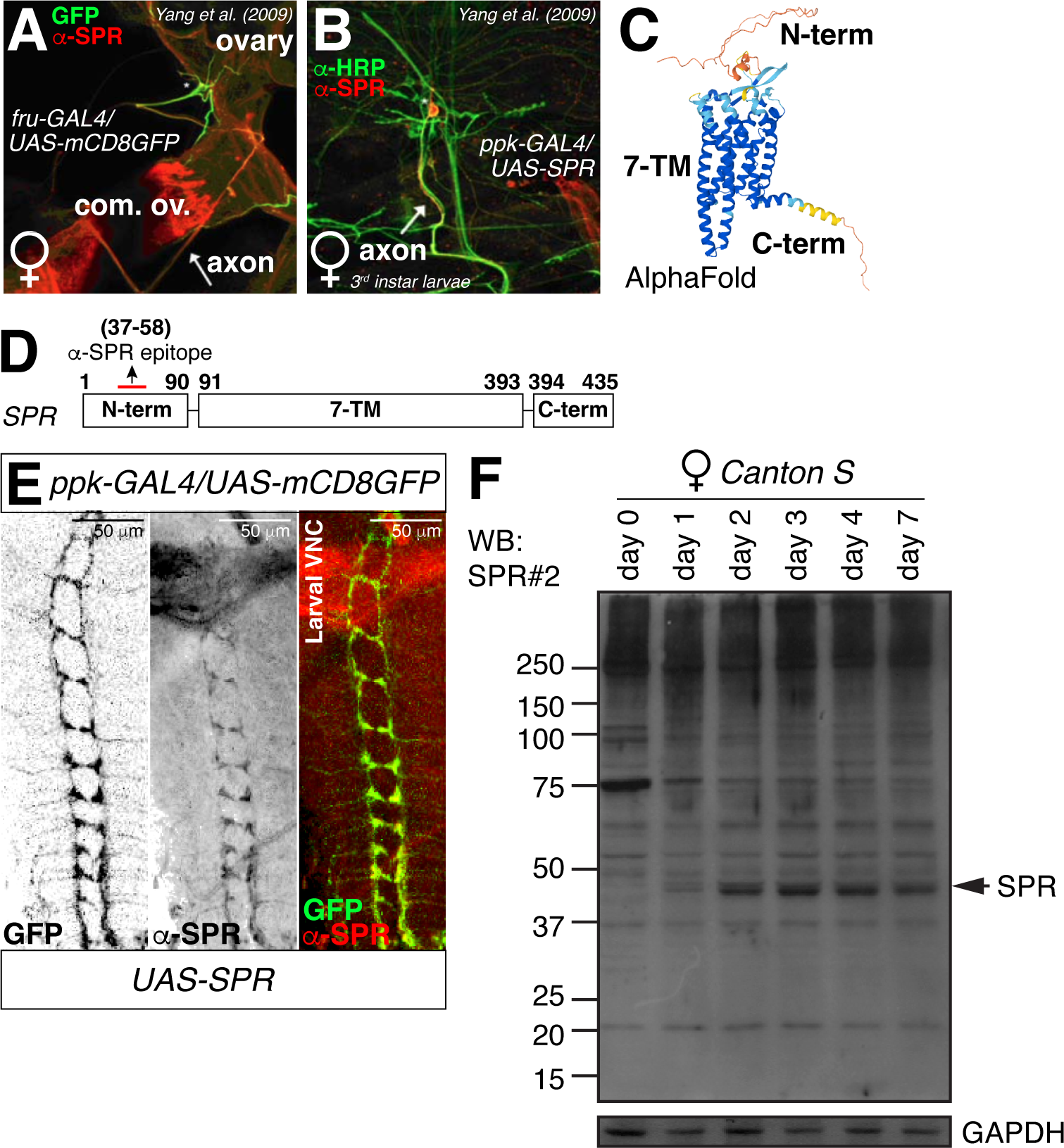
SPR exhibits ectopic expression in the larval body wall da neurons. (A) SPR expression analysis on the lateral oviduct. *fru* neurons were labeled by expression of mCD8-GFP under the control of *fru-GAL4*, and visualized by staining with anti-mCD8 antibodies. SPR was co-stained with anti-SPR antibodies. The overlay shows regions of overlapping mCD8 and SPR staining. The cell body of a *fru/ppk* neuron with its corresponding dendrites is marked by an asterisk and the axon by an arrow. Abbreviation: com. ov., common oviduct. Data were adopted from the study by Yang et al. (2009) (Yang et al. 2009). (B) SPR expression on the uterus was visualized as in Fig. 1A. *fru* neuron cell bodies are marked with arrows. Data were adopted from the study by Yang et al. (2009) (Yang et al. 2009). (C) 3D structure prediction of fruit fly SPR protein contains 7-TM domains generated from AlphaFold. Abbreviations: N-term, N-terminal. C-term, C-terminal. For detailed methods, see the **MATERIALS AND METHODS** for a detailed description of the Protein Domain Predictions procedure used in this study. (D) The linear structure of the SPR protein in *Drosophila melanogaster*, with the numbers indicating the position of each amino acid. The red dash indicates the position where the SPR antibody epitope is recognized. (E) VNC of larva expressing *ppk-GAL4* together with *UAS-mCD8GFP* were immunostained with anti-GFP (green), anti-SPR (red) antibodies. Scale bars represent 50 μm. The left two panels are presented as a grey scale to clearly show the expression patterns of neurons in larval VNC labeled by *ppk-GAL4* driver and anti-SPR. (F) Western blot analysis was performed with anti-SPR antibodies to check the expression level of SPR protein commences post-eclosion into the adult stage in wild type *Drosophila melanogaster*. From lane 1 to lane 6 represent female fruit flies on the 0th, 1st, 2nd, 3rd, 4th, and 7th days after eclosion. The numbers on the left indicate the weight of protein size marker (KDa). For detailed methods, see the **MATERIALS AND METHODS** for a detailed description of the Western Blot procedure used in this study.

## RESULTS

### The ectopic expression of SPR in larval body wall neurons serves as confirmation of its axonal targeting

The fruit fly SPR protein, comprising 435 amino acids, features an N-terminal domain of approximately 90 amino acids, a central segment of roughly 300 amino acids that constitutes seven transmembrane (7-TM) domains, and a C-terminal extension of about 40 amino acids. The confirmation of this molecular structure has been achieved through the application of the advanced AlphaFold computational structural modeling technique which was provided by Fly Base platform (Fig. 1C-D) (Jumper et al. 2021; Gramates et al. 2022; Jenkins et al. 2022; Abramson et al. 2024; Öztürk-Çolak et al. 2024).

To validate the axonal targeting of SPR, we initially elected to employ the larval body wall sensory neuron population, which can be effectively labeled using the *ppk-GAL4* driver line (Grueber et al. 2007). The expression of *UAS-SPR* demonstrated clear colocalization with the axon tracts of the larval ventral nerve cord (VNC), as evidenced by *ppk-GAL4*-driven mCD8GFP fluorescence (Fig. 1E). Given that SPR expression commences post-eclosion into the adult stage, as confirmed by western blot analysis (Fig. 1F), we infer that the observed axonal expression of SPR in the larval VNC is attributable to the artificial expression of the *UAS-SPR* transgene. The selective expression of SPR in the late adult female body was independently validated using previously characterized antibodies (Fig. S1A) (Yapici et al. 2008).

### The N-terminal domain of SPR is essential for its axonal targeting in diverse model systems

To delineate the specific domains of SPR responsible for axonal targeting within the larval VNC, we generated a series of *UAS-SPR* transgenes appended with a C-terminal FLAG tag and established corresponding fly strains (Fig. 2A). Expression of the full-length SPR protein using the *ppk-GAL4* driver confirmed previous observations of SPR targeting to the larval VNC axonal tracts (cf. Fig. 1E with Fig. 2B). Notably, the deletion of the N-terminus in the *UAS-SPRΔN-FLAG*, the C-terminus in the *UAS-SPRΔC-FLAG*, and both termini in the *UAS-SPRΔNC-FLAG* transgenes resulted in a failure to localize to the VNC axonal tracts (Fig. 2C-D and S1B). These findings indicate that both the N- and C-termini of SPR are indispensable for its long-range axonal tract localization.

**Figure 2.**
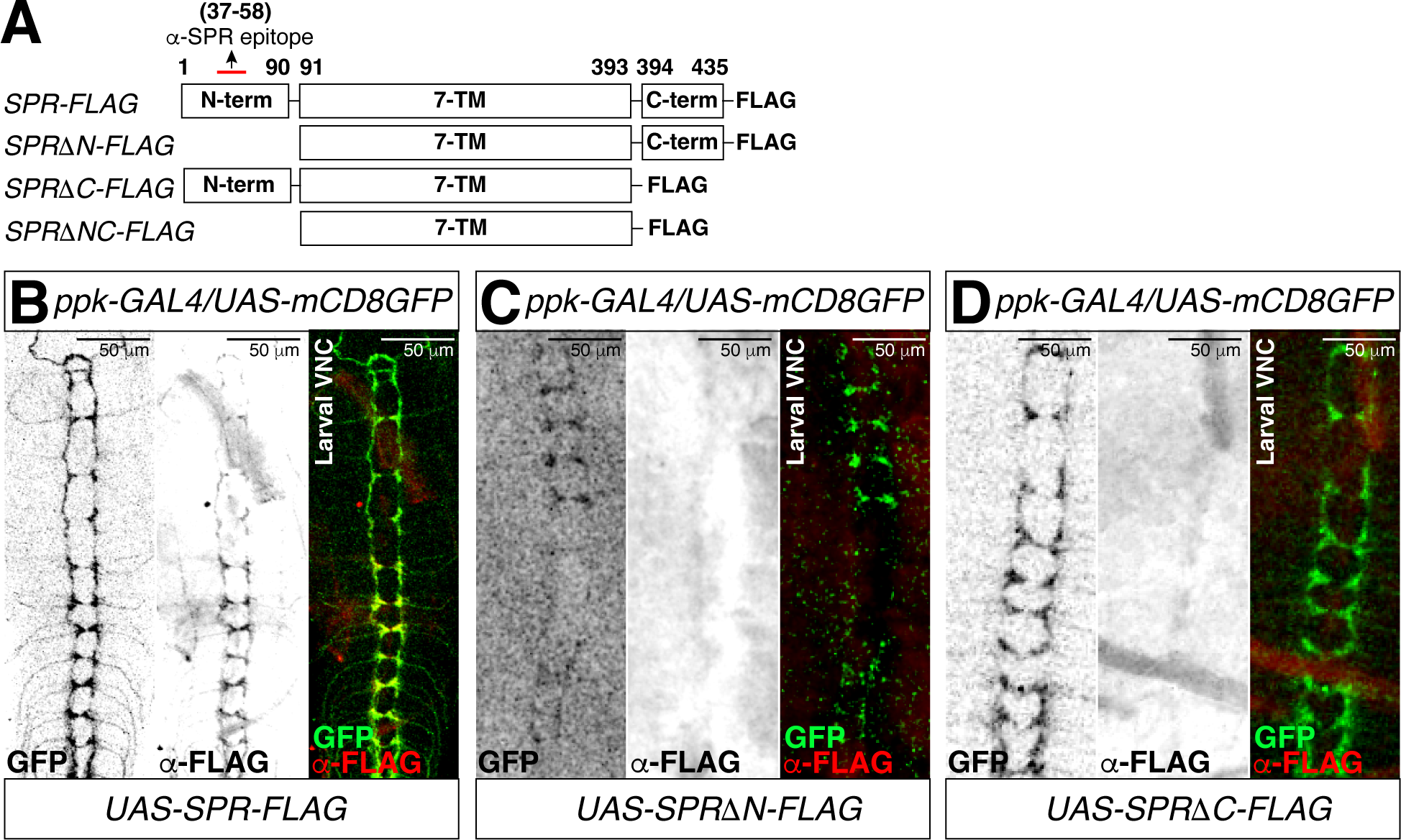
Axonal targeting patterns of SPR in different model systems. (A) The linear structure of a series of SPR protein appended with a C-terminal FLAG tag in *Drosophila melanogaster*, with the numbers indicating the position of each amino acid. (B-D) VNC of larva expressing *ppk-GAL4* together with *UAS-mCD8GFP* were immunostained with anti-GFP (green), anti-FLAG (red) antibodies. ΔN refers to the N-terminal truncation of the SPR protein, ΔC to the C-terminal truncation, and ΔNC to the simultaneous removal of both N- and C-terminal regions of the SPR protein during plasmid construction. Scale bars represent 50 μm. The left two panels are presented as a grey scale to clearly show the expression patterns of neurons in larva VNC labeled by *ppk-GAL4* driver and anti-FLAG.

To investigate the determinants of proximal axonal localization within the SPR protein domains, we utilized dendritic arborization (da) neurons in the larval body wall, which are also amenable to labeling by the *ppk-GAL4* driver (Song et al. 2007; Shimono et al. 2009). The full-length SPR and deletion of the C-terminus of SPR å found to successfully target the axons of da neurons (Fig. 3A and C). However, upon deletion of the N-terminus of SPR, the transgene failed to localize to the axons of da neurons (Fig. 3B and D). These results implicate the N-terminus of SPR as a crucial factor for its proximal axonal localization.

**Figure 3.**
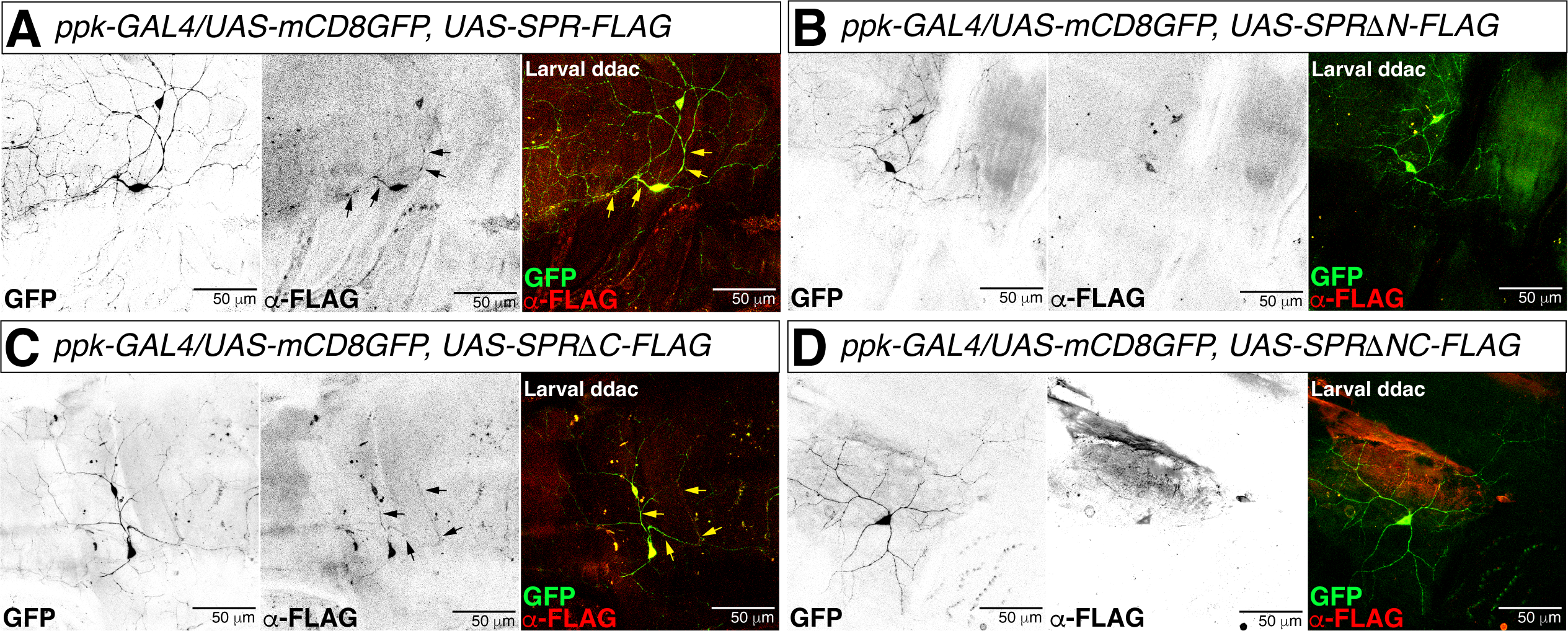
Colocalization patterns of *ppk* and SPR axonal targeting in the larval cuticle of different model systems. (A-D) Body wall of larva expressing *ppk-GAL4* together with *UAS-mCD8GFP* were immunostained with anti-GFP (green), anti-FLAG (red) antibodies. Scale bars represent 50 μm. The left two panels are presented as a grey scale to clearly show the expression patterns of neurons in larval VNC labeled by *ppk-GAL4* driver and anti-FLAG.

Mouse hippocampal neuronal cultures facilitate the mapping of axonal protein localization through a controlled, accessible system that maintains neuronal morphology. This model allows for the manipulation of genetic and environmental factors, enabling the study of axonal targeting mechanisms with high reproducibility and temporal resolution (Cramer and Tyagarajan 2024). In the pursuit of defining neuronal structure, microtubule-associated protein 2 (MAP2) has been established as a crucial identifier for axonal morphology. The preferential localization of MAP2 to the dendrites of hippocampal neurons cultured *in vitro* facilitates the distinction and identification of axonal protein localization (Caceres et al. 1984).

The full-length SPR protein is accurately targeted to the axons of hippocampal neurons (Fig. 4A), indicating that the fly SPR can be correctly localized in mouse hippocampal neurons through evolutionarily conserved mechanisms. In contrast, the N-terminus deleted SPRΔN-GFP variant did not exhibit proper localization in either axons or dendrites (Fig. 4B), implying that the N-terminus of SPR is essential for the targeting of SPR to neural processes. Notably, the C-terminus deleted SPRΔC-GFP variant maintained proper axonal localization (Fig. 4C), suggesting that the C-terminal region of SPR is not a determinant for axonal targeting. The SPRΔNC variant, lacking both N- and C-termini, also failed to localize to axons (Fig. 4D). Quantification of the axonal localization of different SPR domains in hippocampal neurons confirmed the N-terminal region of SPR as a key determinant of axonal targeting (Fig. 4E).

**Figure 4.**
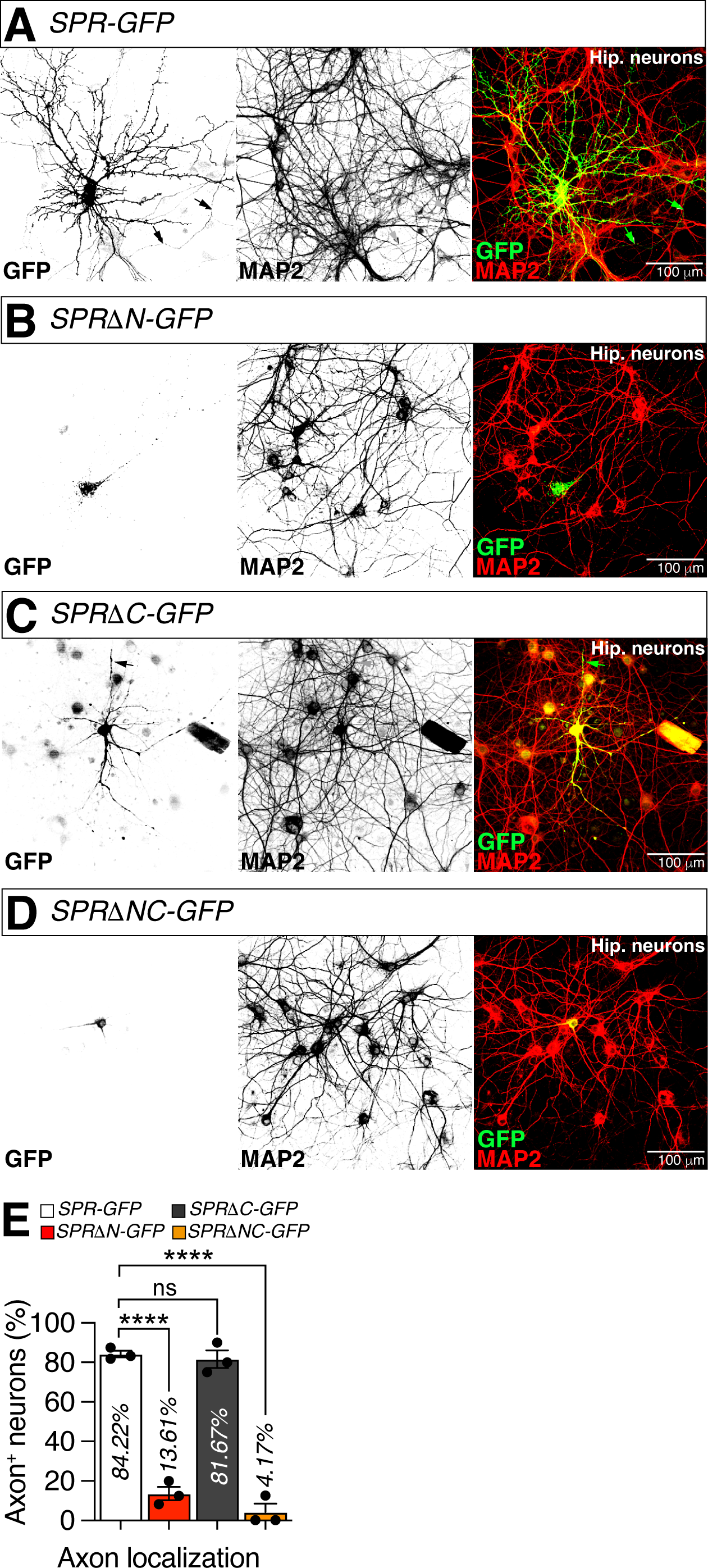
The N-terminal region of SPR as a key determinant of axonal targeting. (A-D) Hippocampal neurons of mouse expressing a series of SPR protein appended with a GFP tag were immunostained with anti-GFP (green), anti-MAP2 (red) antibodies. Black and green arrow indicates axons. Grey arrow indicates empty with MAP2 staining suggesting that it’s axons not dendrites. Scale bars represent 100 μm. The left two panels are presented as a grey scale to clearly show the expression patterns of neurons in hippocampal neurons labeled by anti-GFP and anti-MAP2. (E) Colocalization analysis of GFP and MAP2 staining, normalized to total GFP and MAP2 areas. Bars represent the GFP (Green) proportion which shows non-overlapping with MAP2 staining with error bars representing SEM. Asterisks represent significant differences, as revealed by the Student’s *t* test and ns represents non-significant difference (*\*p<0.05, **p<0.01, ***p< 0.001, ****p< 0.0001*). The same symbols for statistical significance are used in all other figures. See the **MATERIALS AND METHODS** for a detailed description of the colocalization analysis used in this study.

To delineate the essential components of SPR responsible for its axonal localization, we generated a series of SPR transgenes with N-terminal deletions. The deletion of the initial 20 amino acids did not impair axonal targeting (Fig. 5A), implying that these residues are not pivotal for axonal localization. Similarly, the removal of the first 40 amino acids preserved proper axonal localization (Fig. 5B), indicating that these amino acids are also non-essential for this process. Axonal localization of SPR became disrupted upon deletion of the first 60 amino acids (Fig. 5C), suggesting that the region encompassing amino acids 41-60 contains key determinants for axonal targeting. Deletions extending to the first 80 and 90 amino acids completely abrogated axonal localization (Fig. 5D-E), indicating that the critical determinants for axonal targeting of SPR are contained within the first 80 amino acids. Quantitative analysis of axonal localization confirmed that the critical region for axonal targeting is confined to amino acids 41-80 (Fig. 5F).

**Figure 5.**
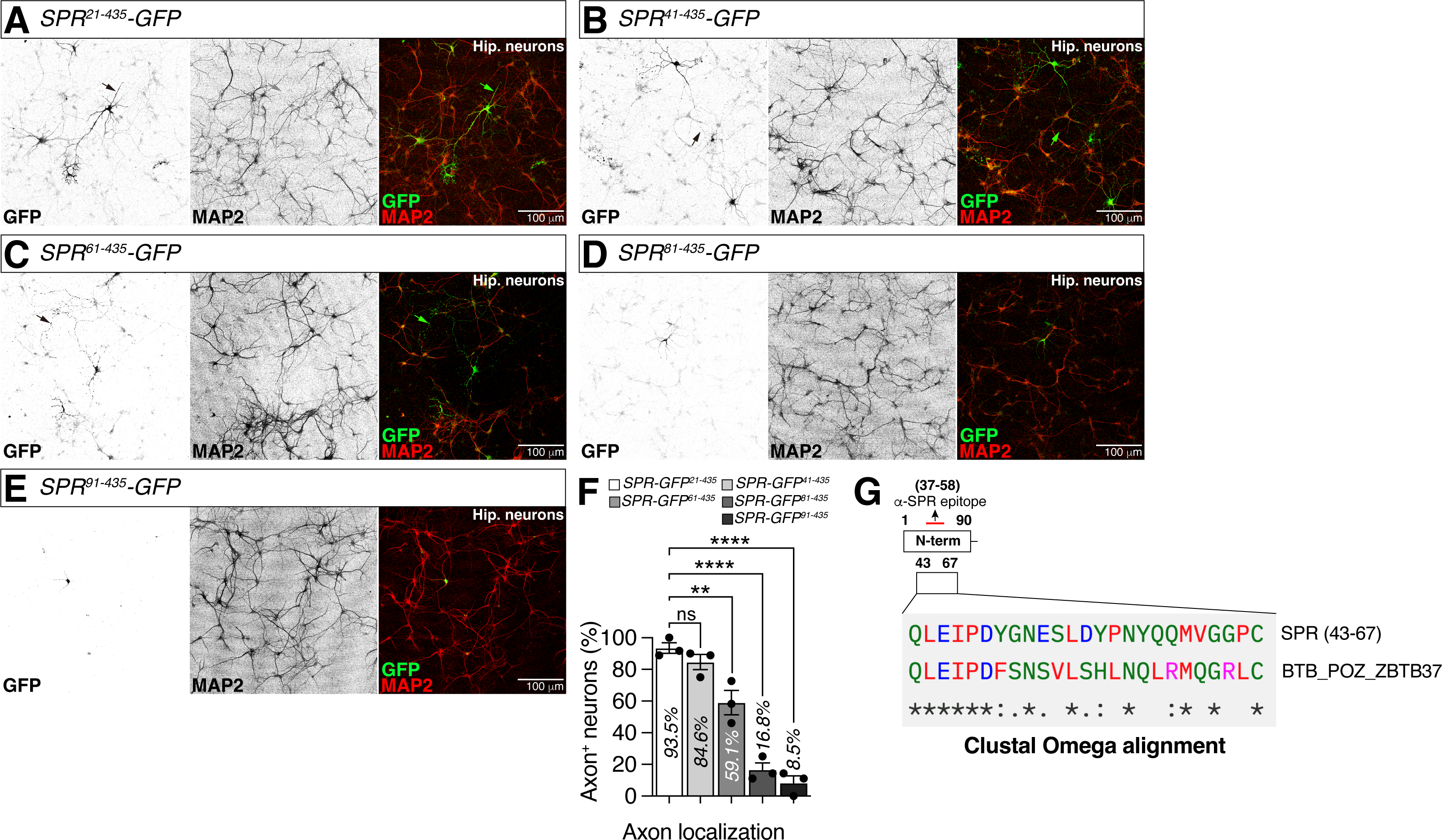
The amino acids 41-80 of SPR is critical region for axonal targeting. (A-E) Hippocampal neurons of mouse expressing SPR protein with a series of N-terminal deletions appended with a GFP tag were immunostained with anti-GFP (green), anti-MAP2 (red) antibodies. Black and green arrow indicates axons. Grey arrow indicates empty with MAP2 staining suggesting that it’s axons not dendrites. Scale bars represent 100 μm. The left two panels are presented as a grey scale to clearly show the expression patterns of neurons in hippocampal neurons labeled by anti-GFP and anti-MAP2. (F) Colocalization analysis of GFP and MAP2 staining, normalized to total GFP and MAP2 areas. (G) Protein sequence alignment analysis of N-terminal amino acid sequence of SPR protein from residues 43 to 67 compared with the BTB/POZ family protein sequences. See the **MATERIALS AND METHODS** for a detailed description of the protein sequence alignment analysis used in this study.

To elucidate the molecular basis of SPR’s axonal localization, we employed the MOTIF Search tool, a bioinformatics resource provided by the Protein Database at the GenomeNet (Kanehisa et al. 2002). This computational approach allowed us to identify potential peptide sequences within the N-terminus of SPR that may serve as axon localization signals. By analyzing the amino acid composition and sequence motifs of the N-terminus, we aimed to uncover conserved patterns or domains that are characteristic of proteins with axonal targeting capabilities. The results of this analysis provided insights into the structural elements that might be responsible for SPR’s specific axonal targeting, thereby facilitating a deeper understanding of the molecular mechanisms underlying axonal localization in neurons.

Through bioinformatics analysis, we identified that the N-terminus of SPR exhibits high similarity to proteins belonging to the Broad-complex, Tramtrack, and Bric-à-brac/poxvirus and zinc finger (BTB/POZ) family (Fig. 5G), which are known for their diverse roles and the presence of a conserved protein-protein interaction domain, the BTB domain (Chaharbakhshi and Jemc 2016; Bonchuk et al. 2023). The homologous region of SPR with the BTB domain spans from amino acids 43 to 67, aligning with the domain mapping results obtained from hippocampal neuronal cultures (Figures 5A-F).

## DISCUSSION

This paper investigates the mechanisms underlying the axonal localization of the SPR in *Drosophila melanogaster*. SPR, a G-protein coupled receptor crucial for female post-mating responses, exhibits axonal targeting in the larval VNC and hippocampal neurons. Our study employed various techniques including transgenic expression, neuronal labeling, and bioinformatics analysis to identify the domains responsible for axonal targeting. Our results reveal that the N-terminal domain of SPR is essential for its axonal localization in both *Drosophila* and mouse models. Deletion of the N-terminus abolished axonal targeting, while the C-terminal domain played a lesser role. Further analysis narrowed down the critical region for axonal targeting to amino acids 41-80 within the N-terminus. Bioinformatics analysis identified a homology between this region and the BTB domain found in proteins with axonal targeting capabilities, suggesting that the BTB domain might be involved in the axonal localization of SPR.

The primary function of SPR lies in orchestrating the post-mating behavioral responses (PMRs) of female *Drosophila melanogaster*. One of the most pronounced effects of PMR is the reduction in female receptivity (Kubli 2003). To assess the impact of axonal localization on this behavioral phenotype, we conducted receptivity assays using virgin and mated females expressing various SPR variants in SPR deficiency background. Consistent with previous reports (Häsemeyer et al. 2009; Yang et al. 2009), females with the SPR deficiency *Df^exel6234^* exhibited a robust remating phenotype. However, intriguingly, expression of the full-length SPR in fru-positive neurons fully rescued this remating phenotype (groups 1 and 2 in Fig. S1C). Even more surprising, expression of the N-terminus-deleted SPR in fru-positive neurons also fully rescued the remating phenotype, albeit at a level comparable to that of the full-length SPR (group 3 in Fig. S1C). In contrast, expression of the C-terminus-deleted or N- and C-terminus-deleted SPR variants in fru-positive neurons failed to rescue the remating phenotype of SPR mutant females (groups 4 and 5 in Fig. S1C). These findings suggest that the N-terminus of SPR is not a critical determinant for the remating phenotype of females. Consequently, we conclude that axonal localization of SPR is not essential for regulating female remating behavior.

The identification of a BTB domain-like region in the N-terminus of SPR raises intriguing possibilities regarding its role in axonal targeting and function. The BTB domain is a well-characterized protein-protein interaction domain found in various proteins involved in diverse cellular processes, including transcriptional regulation, chromatin remodeling, and protein degradation (Chaharbakhshi and Jemc 2016; Bonchuk et al. 2023). Its presence in SPR suggests that this domain might mediate protein-protein interactions crucial for its axonal localization and function.

One potential mechanism by which the BTB domain-like region could contribute to axonal targeting involves interactions with cytoskeletal proteins or motor proteins. For instance, BTB proteins have been shown to interact with microtubule-associated proteins (MAPs), which regulate microtubule dynamics and organization (Ding et al. 2002). This interaction could facilitate the transport of SPR along axonal microtubules using motor proteins, ensuring its delivery to the appropriate axonal compartments. Additionally, BTB proteins have been implicated in interactions with membrane proteins, potentially facilitating the insertion of SPR into the axonal plasma membrane (Chaharbakhshi and Jemc 2016; Bonchuk et al. 2023).

Furthermore, the BTB domain-like region might play a role in the formation of multi-protein complexes involving SPR. These complexes could mediate the downstream signaling pathways activated by SPR, ultimately influencing post-mating behaviors. For instance, BTB proteins often function as transcriptional regulators, and the BTB domain-like region in SPR could interact with other transcription factors or co-regulators, modulating gene expression in response to sex peptide signaling.

While the precise mechanisms by which the BTB domain-like region contributes to SPR localization and function remain to be elucidated, its identification provides valuable insights into the molecular underpinnings of axonal targeting and post-mating responses in *Drosophila*. Future research could explore the interaction partners of the BTB domain-like region in SPR, investigate its role in protein trafficking and signaling, and determine its contribution to the behavioral phenotypes associated with SPR mutations.

## METHODS

### Fly Stocks and Husbandry

*Drosophila melanogaster* were raised on cornmeal-yeast medium at similar densities to yield adults with similar body sizes. Flies were kept in 12 h light: 12 h dark cycles (LD) at 25 (ZT 0 is the beginning of the light phase, ZT12 beginning of the dark phase). Wild-type flies were Canton-S. Following lines used in this study, *Canton-S* (#64349)*, fru-GAL4* (#66696)*, UAS-mCD8GFP* (#79626), and *ppk-GAL4* (#32079) were obtained from the Bloominton *Drosophila* Stock Center at Indiana University. The following line, *UAS-SPR* was a gift from Barry Dickson (Queensland Brain Institute, Austrailia).

### Immunostaining

The dissection and immunostaining protocols for the experiments are described elsewhere (Kim et al. 2013). After 5 days of eclosion, the *Drosophila* brain was taken from adult flies and fixed in 4% formaldehyde at room temperature for 30 minutes. The sample was washed three times (5 minutes each) in 1% PBT and then blocked in 5% normal goat serum for 30 minutes. The sample next be incubated overnight at 4 with primary antibodies in 1% PBT, followed by the addition of fluorophore-conjugated secondary antibodies for one hour at room temperature. The brain was mounted on plates with an antifade mounting solution (Solarbio) for imaging purposes. Antibodies were used at the following dilutions: Chicken anti-GFP (Invitrogen, 1:500), Rabbit anti-SPR (Customized, 1:500), Alexa-488 donkey anti-chicken (Jackson ImmunoResearch, 1:200), Alexa-555 goat anti-rabbit (Invitrogen, 1:200), mouse anti-FALG (Merck, 1:400), Chicken anti-MAP2 (Invitrogen, 1:2000).

### Protein Domain Predictions

To predict the N-terminal peptide sequence of SPR, MOTIF search tool on GenomeNet (MotifFinder https://www.genome.jp/tools-bin/search_motif_lib) were used in this study. By analyzing the amino acids of the translated gene sequence, we identified potential N-terminal motifs and cross-referenced these predictions with existing biological databases and literature to ensure their relevance and accuracy.

### Protein Sequence Alignment

To perform protein sequence alignment, we utilized the Clustal Omega tool on the EMBL-EBI website (https://www.ebi.ac.uk/Tools/msa/clustalo/). We input the protein sequences of SPR, along with BTB_POZ_ZBTB37 sequences, to conduct the alignment. By examining the aligned sequences, we identified regions of conservation and variation, which were then cross-referenced with existing biological databases and literature to ensure the functional context of the observed patterns.

### Antibody generation

To generate specific antibodies against the SPR protein, we focused on the PTNESQLEIPDYGNESLDYPNC peptide region. This peptide was synthesized and used to immunize rabbits, following standard protocols for polyclonal antibody production. The resulting antiserum was then purified using affinity chromatography to obtain anti-rabbit polyclonal antibodies specific to the PTNESQLEIPDYGNESLDYPNC region of SPR. The specificity and titer of the antibodies were validated Western blotting against the recombinant SPR protein and control samples.

### Molecular Cloning

For the generation of plasmid expressing SPR and GFP under the control of the SPR activation sequence, we cloned the SPR gene sequence into two different vectors: the pUAST-attB vector and the pCMV-GFP vector, which tags the protein at its C-terminus with GFP (https://www.addgene.org/11153/). To create a FLAG-tagged version of SPR, we designed the reverse primer to include the FLAG epitope sequence, as described by Grozdanov (Grozdanov and MacDonald 2015) (https://www.ptglab.com/news/blog/flag-tag-and-3x-flag-tag/). This primer was used in a PCR reaction to amplify the SPR sequence with the FLAG tag integrated. The PCR product was then cloned into the UAS-SPR-FLAG vector, and the construct was transformed into fly embryos to generate transgenic lines. The successful integration and expression of the FLAG-tagged SPR in the plasmid were verified through PCR and immunohistochemical analyses using anti-FLAG antibodies.

### Receptivity Assay

All flies were aged for 4–5 days post-eclosion before being subjected to behavioral analysis. The females and males were kept in groups of 3–4 individuals in small containers. To evaluate female receptivity, a single female was introduced into a small (1 cm x 1 cm) chamber lined with grape agar, accompanied by two naive Canton S male flies. The female was deemed receptive if she engaged in mating within a 20-minute timeframe (Häsemeyer et al. 2009; Yang et al. 2009).

### Western Blot Analysis

The similar method was described elsewhere (Kim et al. 2007). For Western blotting, flies were homogenized in pools with cold NP-40 lysis buffer with a motorized pestle in 1.5 mL microcentrifuge tubes. Homogenates were transferred to fresh 1.5ml microfuge tubes, and 10 μL of homogenized sample was used for protein quantification. Finally, the precipitated proteins were resolved by SDS-PAGE and transferred to a nitrocellulose or PVDF membrane for subsequent detection with the appropriate secondary antibodies and visualization using an enhanced chemiluminescence system.

### Larval Body Wall Neurons Live Imaging

For live imaging of larval body wall neurons, the flies were reared at 25 °C in density-controlled vials. Third instar larvae at 96 h after egg laying (AEL) were mounted in halocarbon oil and confocal images of ddaC dendrites were collected with a Leica SP5 laser scanning microscope (Han et al. 2011).

### Hippocampal Neuronal Culture and Imaging

To culture hippocampal neurons, the wild type mice brain is carefully extracted from newborn mice, and the hippocampus is dissected and dissociated into individual cells. These cells are then plated onto coated coverslips and maintained in culture. Later, lipofectine are used to introduce specific DNA plasmid into the neurons. Finally, the neurons are stimulated to induce dendritic plasticity, and the properties of the dendrites are measured using confocal microscopy techniques (Caceres et al. 1984; Shi et al. 2003; Cramer and Tyagarajan 2024).

### Quantification of Axonal Localization

To perform colocalization analysis of multi-color fluorescence microscopy images in this study, we employed ImageJ software (Daly.). We merged image channels to form a composite with accurate color representation and applied a threshold to isolate yellow pixels, signifying colocalization. The measured “area” values represented the colocalization zones between fluorophores. To determine the colocalization percentage relative to the total area of interest (e.g., GFP or RFP), we adjusted thresholds to capture the full fluorophore areas and remeasured to obtain total areas. The colocalization efficiency was calculated by dividing the colocalized area by the total fluorophore area. All samples were imaged uniformly.

### Statistical Tests

Two-sided Student’s t tests was used in this study. The mean ± standard error (s.e.m) (*\*\*\*\* = p < 0.0001, *** = p < 0.001, ** = p < 0.01, * = p < 0.05*). All analysis was done in GraphPad (Prism). Individual tests and significance are detailed in figure legends.

## ACKNOWLEDGEMENTS

We extend our gratitude to the individuals who provided us with the *Drosophila melanogaster* strains and research reagents. Specifically, we acknowledge Dr. Barry Dickson (Queensland Brain Institute, Australia) for supplying the UAS-SPR fly strain and the anti-SPR antibody, which was instrumental in validating the specificity of our developed anti-SPR antibody. We are also thankful to Drs. Yuh Nung Jan and Lily Yeh Jan (University of California San Francisco, USA) for their invaluable guidance and ongoing support throughout this project. This research was supported by Startup funds from HIT Center for Life Science (HCLS) to WJK.

---

**Figure S1. The axonal localization of SPR is not essential for regulating female remating behavior.**

(A) Western blot analysis was performed with anti-SPR antibodies to check the expression level of SPR protein commences post-eclosion into the adult stage in wild type *Drosophila melanogaster*. From lane 1 to lane 6 represent female fruit flies on the 0th, 1st, 2nd, 3rd, 4th, and 7th days after eclosion.

(B) VNC of larva expressing *ppk-GAL4* together with *UAS-mCD8GFP, and UAS-SPRΔ*NC-FLAG were immunostained with anti-GFP (green), anti-FLAG (red) antibodies. Scale bars represent 50 μm. The left two panels are presented as a grey scale to clearly show the expression patterns of neurons in larval VNC labeled by *ppk-GAL4* driver and anti-FLAG.

(C) Receptivity assays for virgin and mated females expressing various *UAS-SPR* variants in SPR deficiency background and *fru*-positive neurons. Asterisks represent significant differences, as revealed by the Student’s t test and ns represents non-significant difference (*p<0.05, **p<0.01, ***p< 0.001, ****p< 0.0001). See the MATERIALS AND METHODS for a detailed description of the receptivity assays used in this study.

## REFERENCES

Abramson J, Adler J, Dunger J, Evans R, Green T, Pritzel A, Ronneberger O, Willmore L, Ballard AJ, Bambrick J, et al. 2024. Accurate structure prediction of biomolecular interactions with AlphaFold 3. Nature. 630(8016):493–500. doi:10.1038/s41586-024-07487-w.

Audsley N, Down RE. 2015. G protein coupled receptors as targets for next generation pesticides. Insect Biochem Molec. 67:27–37. doi:10.1016/j.ibmb.2015.07.014.

Avila FW, Mattei AL, Wolfner MF. 2015. Sex peptide receptor is required for the release of stored sperm by mated Drosophila melanogaster females. J Insect Physiol. 76:1–6. doi:10.1016/j.jinsphys.2015.03.006.

Boivin B, Vaniotis G, Allen BG, Hébert TE. 2008. G Protein-Coupled Receptors in and on the Cell Nucleus: A New Signaling Paradigm? J Recept Signal Transduct. 28(1–2):15–28. doi:10.1080/10799890801941889.

Bonchuk A, Balagurov K, Georgiev P. 2023. BTB domains: A structural view of evolution, multimerization, and protein–protein interactions. BioEssays. 45(2):e2200179. doi:10.1002/bies.202200179.

Caceres A, Banker G, Steward O, Binder L, Payne M. 1984. MAP2 is localized to the dendrites of hippocampal neurons which develop in culture. Dev Brain Res. 13(2):314–318. doi:10.1016/0165-3806(84)90167-6.

Caers J, Verlinden H, Zels S, Vandersmissen HP, Vuerinckx K, Schoofs L. 2012. More than two decades of research on insect neuropeptide GPCRs: an overview. Front Endocrinol. 3:151. doi:10.3389/fendo.2012.00151.

Chaharbakhshi E, Jemc JC. 2016. Broad complex, tramtrack, and bric à brac (BTB) proteins: Critical regulators of development. genesis. 54(10):505–518. doi:10.1002/dvg.22964.

Chapman T, Bangham J, Vinti G, Seifried B, Lung O, Wolfner MF, Smith HK, Partridge L. 2003. The sex peptide of Drosophila melanogaster: Female post-mating responses analyzed by using RNA interference. Proc National Acad Sci. 100(17):9923–9928. doi:10.1073/pnas.1631635100.

Cramer TML, Tyagarajan SK. 2024. Protocol for the culturing of primary hippocampal mouse neurons for functional in vitro studies. STAR Protoc. 5(2):102991. doi:10.1016/j.xpro.2024.102991.

Daly. C. Colocalisation tutorial using ImageJ [Video]. YouTube. https://www.youtube.com/watch?v=rRyJnFo57xU.

Ding J, Liu J-J, Kowal AS, Nardine T, Bhattacharya P, Lee A, Yang Y. 2002. Microtubule-associated protein 1B. J Cell Biol. 158(3):427–433. doi:10.1083/jcb.200202055.

Feng K, Palfreyman MT, Häsemeyer M, Talsma A, Dickson BJ. 2014. Ascending SAG Neurons Control Sexual Receptivity of Drosophila Females. Neuron. 83(1):135–148. doi:10.1016/j.neuron.2014.05.017.

Gramates LS, Agapite J, Attrill H, Calvi BR, Crosby MA, Santos Gilberto dos, Goodman JL, Goutte-Gattat D, Jenkins VK, Kaufman T, et al. 2022. FlyBase: a guided tour of highlighted features. Genetics. 220(4):iyac035. doi:10.1093/genetics/iyac035.

Grozdanov PN, MacDonald CC. 2015. Generation of Plasmid Vectors Expressing FLAG-tagged Proteins Under the Regulation of Human Elongation Factor-1α Promoter Using Gibson Assembly. J Vis Exp.(96). doi:10.3791/52235.

Grueber WB, Ye B, Yang C-H, Younger S, Borden K, Jan LY, Jan Y-N. 2007. Projections of Drosophila multidendritic neurons in the central nervous system: links with peripheral dendrite morphology. Development. 134(1):55–64. doi:10.1242/dev.02666.

Han C, Jan LY, Jan Y-N. 2011. Enhancer-driven membrane markers for analysis of nonautonomous mechanisms reveal neuron–glia interactions in Drosophila. Proc National Acad Sci. 108(23):9673–9678. doi:10.1073/pnas.1106386108.

Hanlon CD, Andrew DJ. 2015. Outside-in signaling – a brief review of GPCR signaling with a focus on the Drosophila GPCR family. J Cell Sci. 128(19):3533–3542. doi:10.1242/jcs.175158.

Häsemeyer M, Yapici N, Heberlein U, Dickson BJ. 2009. Sensory Neurons in the Drosophila Genital Tract Regulate Female Reproductive Behavior. Neuron. 61(4):511–518. doi:10.1016/j.neuron.2009.01.009.

Hauser F, Cazzamali G, Williamson M, Blenau W, Grimmelikhuijzen CJP. 2006. A review of neurohormone GPCRs present in the fruitfly Drosophila melanogaster and the honey bee Apis mellifera. Prog Neurobiol. 80(1):1–19. doi:10.1016/j.pneurobio.2006.07.005.

Hussain A, Üçpunar HK, Zhang M, Loschek LF, Kadow ICG. 2016. Neuropeptides Modulate Female Chemosensory Processing upon Mating in Drosophila. PLoS Biol. 14(5):e1002455. doi:10.1371/journal.pbio.1002455.

Jenkins VK, Larkin A, Thurmond J, Consortium F. 2022. Using FlyBase: A Database of Drosophila Genes and Genetics. Methods Mol Biol (Clifton, NJ). 2540:1–34. doi:10.1007/978-1-0716-2541-5_1.

Jumper J, Evans R, Pritzel A, Green T, Figurnov M, Ronneberger O, Tunyasuvunakool K, Bates R, Žídek A, Potapenko A, et al. 2021. Highly accurate protein structure prediction with AlphaFold. Nature. 596(7873):583–589. doi:10.1038/s41586-021-03819-2.

Kanehisa M, Goto S, Kawashima S, Nakaya A. 2002. The KEGG databases at GenomeNet. Nucleic Acids Res. 30(1):42–46. doi:10.1093/nar/30.1.42.

Kim WJ, Jan LY, Jan YN. 2013. A PDF/NPF Neuropeptide Signaling Circuitry of Male Drosophila melanogaster Controls Rival-Induced Prolonged Mating. Neuron. 80(5):1190–1205. doi:10.1016/j.neuron.2013.09.034.

Kim WJ, Kim JH, Jang SK. 2007. Anti-inflammatory lipid mediator 15d-PGJ2 inhibits translation through inactivation of eIF4A. The EMBO journal. 26(24):5020–5032.

Kubli E. 2003. Sex-peptides: seminal peptides of the Drosophila male. Cell Mol Life Sci Cmls. 60(8):1689–1704. doi:10.1007/s00018-003-3052.

Lasiecka ZM, Yap CC, Vakulenko M, Winckler B. 2008. Chapter 7 Compartmentalizing the Neuronal Plasma Membrane From Axon Initial Segments to Synapses. Int Rev Cell Mol Biol. 272(J. Cell Biol.1522001):303–389. doi:10.1016/s1937-6448(08)01607-9.

Oh Y, Yoon S-E, Zhang Q, Chae H-S, Daubnerová I, Shafer OT, Choe J, Kim Y-J. 2014. A Homeostatic Sleep-Stabilizing Pathway in Drosophila Composed of the Sex Peptide Receptor and Its Ligand, the Myoinhibitory Peptide. Plos Biol. 12(10):e1001974. doi:10.1371/journal.pbio.1001974.

Öztürk-Çolak A, Marygold SJ, Antonazzo G, Attrill H, Goutte-Gattat D, Jenkins VK, Matthews BB, Millburn G, Santos G dos, Tabone CJ, et al. 2024. FlyBase: updates to the Drosophila genes and genomes database. GENETICS. 227(1):iyad211. doi:10.1093/genetics/iyad211.

Rezával C, Nojima T, Neville MC, Lin AC, Goodwin SF. 2014. Sexually Dimorphic Octopaminergic Neurons Modulate Female Postmating Behaviors in Drosophila. Curr Biol. 24(7):725–730. doi:10.1016/j.cub.2013.12.051.

Shi S-H, Jan LY, Jan Y-N. 2003. Hippocampal Neuronal Polarity Specified by Spatially Localized mPar3/mPar6 and PI 3-Kinase Activity. Cell. 112(1):63–75. doi:10.1016/s0092-8674(02)01249-7.

Shimono K, Fujimoto A, Tsuyama T, Yamamoto-Kochi M, Sato M, Hattori Y, Sugimura K, Usui T, Kimura K, Uemura T. 2009. Multidendritic sensory neurons in the adult Drosophila abdomen: origins, dendritic morphology, and segment- and age-dependent programmed cell death. Neural Dev. 4(1):37. doi:10.1186/1749-8104-4-37.

Song W, Onishi M, Jan LY, Jan YN. 2007. Peripheral multidendritic sensory neurons are necessary for rhythmic locomotion behavior in Drosophila larvae. Proc National Acad Sci. 104(12):5199–5204. doi:10.1073/pnas.0700895104.

Stefan CJ, Overton MC, Blumer KJ. 1998. Mechanisms Governing the Activation and Trafficking of Yeast G Protein-coupled Receptors. Mol Biol Cell. 9(4):885–899. doi:10.1091/mbc.9.4.885.

Tsuda M, Peyre J-B, Asano T, Aigaki T. 2015. Visualizing Molecular Functions and Cross-Species Activity of Sex-Peptide in Drosophila. Genetics. 200(4):1161–1169. doi:10.1534/genetics.115.177550.

Vandersmissen HP, Nachman RJ, Broeck JV. 2013. Sex peptides and MIPs can activate the same G protein-coupled receptor. Gen Comp Endocr. 188:137–143. doi:10.1016/j.ygcen.2013.02.014.

Wolfner MF. 2002. The gifts that keep on giving: physiological functions and evolutionary dynamics of male seminal proteins in Drosophila. Heredity. 88(2):85–93. doi:10.1038/sj.hdy.6800017.

Yang C, Rumpf S, Xiang Y, Gordon MD, Song W, Jan LY, Jan Y-N. 2009. Control of the Postmating Behavioral Switch in Drosophila Females by Internal Sensory Neurons. Neuron. 61(4):519–526. doi:10.1016/j.neuron.2008.12.021.

Yang YT, Hu SW, Li X, Sun Y, He P, Kohlmeier KA, Zhu Y. 2023. Sex peptide regulates female receptivity through serotoninergic neurons in Drosophila. iScience. 26(3):106123. doi:10.1016/j.isci.2023.106123.

Yapici N, Kim Y-J, Ribeiro C, Dickson BJ. 2008. A receptor that mediates the post-mating switch in Drosophila reproductive behaviour. Nature. 451(7174):33–37. doi:10.1038/nature06483.

