## Supplementary figures and images for "Investigating the role of axonal localization of sex peptide receptor in post-mating responses of female *Drosophila melanogaster*"

### Supplemental File 1

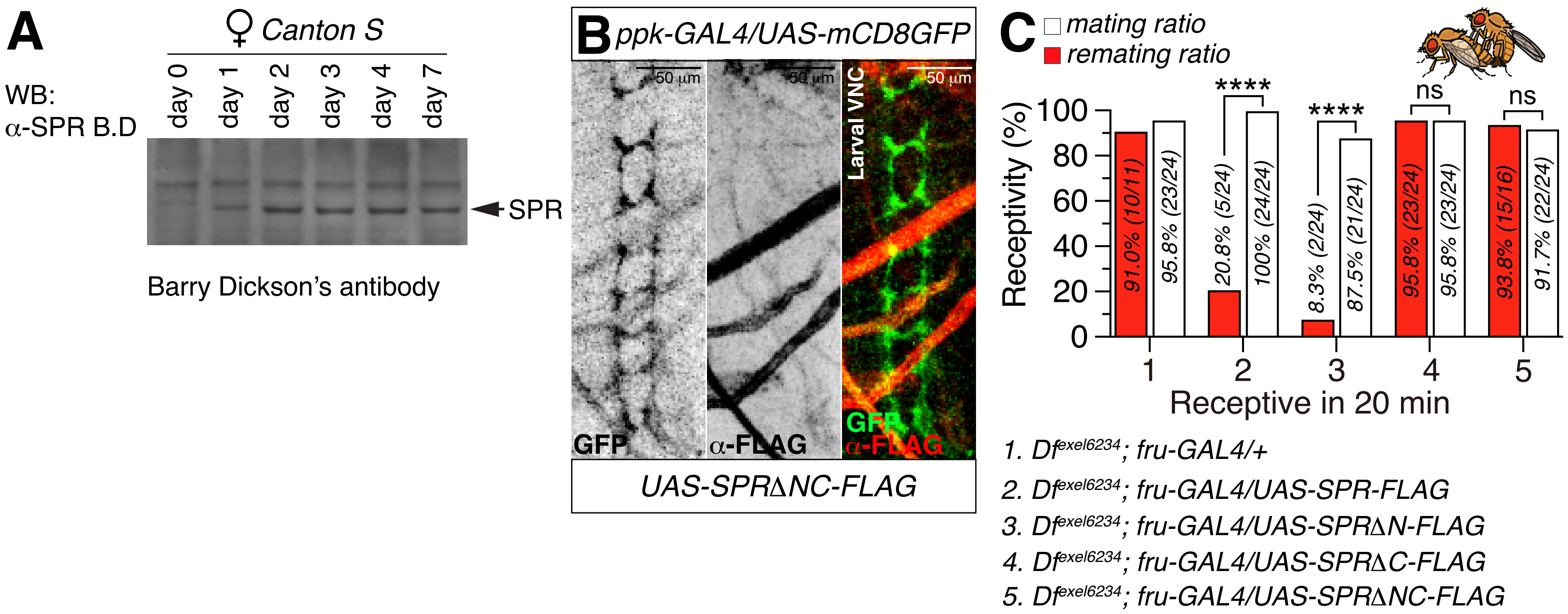
